# Efficient horizontal transmission without viral super-spreaders may cause the high prevalence of STLV-1 infection in Japanese macaques

**DOI:** 10.1101/639062

**Authors:** Megumi Murata, Jun-ichirou Yasunaga, Ayaka Washizaki, Yohei Seki, Wei Keat Tan, Takuo Mizukami, Masao Matsuoka, Hirofumi Akari

## Abstract

Simian T-cell leukemia virus type-1 (STLV-1) is disseminated among various non-human primate species and is closely related to human T-cell leukemia virus type-1 (HTLV-1), the causative agent of adult T-cell leukemia and HTLV-1-associated myelopathy/tropical spastic paraparesis. Notably, the prevalence of STLV-1 infection in Japanese macaques (JMs) is estimated to be much greater than that in other non-human primates; however, the mechanism and mode of STLV-1 transmission remain unknown. We hypothesized that a substantial proportion of infected macaques may play a critical role as viral super-spreaders for efficient inter-individual transmission leading to the high prevalence of infection. To address this, we examined a cohort of 280 JMs reared in a free-range facility for levels of anti-STLV-1 antibody titers (ABTs) and STLV-1 proviral loads (PVLs). We found that the prevalence of STLV-1 in the cohort reached up to 65% (180/280), however, the ABTs and PVLs were normally distributed with mean values of 4076 and 0.62%, respectively, which were comparable to those of HTLV-1-infected humans. Contrary to our expectations, we did not observe the macaques with abnormally high PVLs and poor ABTs, and therefore, the possibility of viral super-spreaders was unlikely. Results from further analyses regarding age-dependent changes in STLV-1 prevalence and a longitudinal follow-up of STLV-1 seroconversion strongly suggest that frequent horizontal transmission is a major route of STLV-1 infection, probably due to the unique social ecology of JMs associated with environmental adaptation.

**Importance:** We investigated the cause of the high prevalence of STLV-1 infection in the studied JMs cohort. Contrary to our expectations, the potential viral super-spreaders as shown by abnormally high PVLs and poor ABTs were not observed among the JMs. Rather, the ABTs and PVLs among the infected JMs were comparable to those of HTLV-1-infected humans although the prevalence of HTLV-1 in humans is much less than the macaques. Further analyses demonstrate that the prevalence drastically increased over one year of age and most of these animals over 6 years of age were infected with STLV-1, and that in the longitudinal follow-up study frequent seroconversion occurred in not only infants but also in juvenile and adult seronegative monkeys (around 20% per year). This is the first report showing that frequent horizontal transmission without viral super-spreaders may cause high prevalence of STLV-1 infection in JMs.

## Introduction

Simian T-cell leukemia viruses (STLVs) are classified into the Deltaretrovirus genus, which includes human T-cell leukemia viruses (HTLVs). The first human retrovirus, HTLV-1, was identified in 1980 (1–3), even though the disease entity of adult T-cell leukemia (ATL) had been described in Japan before the identification of this virus (4). Eventually, HTLV-1 was found to be the causative agent of not only ATL but also HTLV-1-associated myelopathy (HAM)/tropical spastic paraparesis (TSP) (1, 2, 5-10). It is estimated that 10–20 million people worldwide are infected with HTLV-1 (11). HTLV-1 infections are endemic in southern Japan, Africa, the Caribbean, Central and South America, and intertropical Africa (12–14). An estimated one million people in Japan are thought to be HTLV-1 carriers, corresponding to 1% of the total population (14–16). In most cases, HTLV-1 infection remains asymptomatic, whereas 5% of carriers develop ATL and/or HAM/TSP (17–24). STLVs infect a variety of non-human primates in Asia and Africa but not in America (25–28). STLV-1 and STLV-2 have human counterparts, HTLV-1 and HTLV-2 (29–32). A third subspecies, STLV-3, was isolated from an Eritrean sacred baboon (*Papio hamadryas*) and a red-capped mangabey (*Cercocebus torquatus*) (33, 34). A recent report showed that STLV-4 was isolated from gorillas and that the virus was endemic to gorillas (35). It has been reported that STLVs are also associated with leukemia/lymphoma (36–40) and that hunting and severe bites by non-human primates are the likely routes of zoonotic transmission of STLVs (26, 41-45).

Japanese macaques (JMs: *Macaca fuscata*) inhabit much of Japan (except Hokkaido and Okinawa). JMs are found infected with STLV-1, and their seroprevalence is much greater than that of other primates (46–52). Watanabe et al. reported that the sequence homology of STLV-1 to that of HTLV-1 was 90% (29). Given this genetic similarity, it was suspected that zoonotic STLV transmission might be, at least in part, the cause of HTLV-1 dissemination among Japanese people. However, phylogenetic analysis between HTLV-1 isolated from Japanese people and STLV-1 isolated from JMs demonstrated that STLV-1 was distinct from HTLV-1 (53). Furthermore, some groups have reported that the geographical distribution of HTLV-1 in Japan did not correspond to the habitat of JMs (50, 54). From genomic and epidemiological evidence, it was concluded that Japanese HTLV-1 originated from Mongoloid people moving from North Asia but not from JM STLV-1 (53, 55).

A high proportion (60% on average) of JMs has been reported as infected with STLV-1, whereas the prevalence of STLV in other natural hosts among non-human primates, including Asian macaques, is generally much lower than with JMs (25, 50-52, 56-62). The reason of the high prevalence remains unknown. However, it was proposed that STLV-1 in JMs may have an alternative transmission route via maternal infection (51). We hypothesized that the substantial proportion of infected macaques may play critical roles as viral super-spreaders for efficient inter-individual transmission, likely due to abnormally high proviral loads (PVLs) and eventual incidence of poor humoral immune response against STLV-1. We recently experienced an outbreak of infectious malignant thrombocytopenia in JMs by simian retrovirus type 4 (SRV-4) infection (63). Importantly, some of the monkeys who developed persistent SRV-4 infection exhibited viremia without an SRV-4-specific antibody response and became viral super-spreaders (64). Taking this example into account, we evaluated antibody titers (ABTs) against STLV-1 and PVLs in the JM cohort.

## Results

To validate the STLV-1 prevalence in JMs, we first examined the anti-STLV-1 ABTs from the plasma of 280 JMs derived from five independent troops originating from inhabitants of different areas. We found that 180 macaques (65%) were seropositive (Table 1), which was generally consistent with previous reports (47, 48, 50, 52). We then determined the variation in the seroprevalence among the troops. The numbers of seropositive individuals were 59, 17, 36, 34, and 34, with a frequency of 68%, 55%, 63%, 56%, and 77%, respectively (Table 1). The seroprevalence was generally comparable with that in wild JMs as previously reported (50, 52). In addition, the rearing density in each troop was not correlated with the seroprevalence, suggesting that relatively higher population density may not cause the high prevalence (Table 1).

**Table 1:** Seroprevalence in Japanese macaques and other parameters having the possibilities of affecting seroprevalence in each area.

|  | Troop A | Troop B | Troop C | Troop D | Troop E | total |
| --- | --- | --- | --- | --- | --- | --- |
| Number of individuals (male/female) | 87(32/55) | 31(9/22) | 57(24/33) | 61(18/43) | 44(19/25) | 280(102/178) |
| STLV-1 seroprevalence |  |  |  |  |  |  |
| Number of positive individuals | 59 | 17 | 36 | 34 | 34 | 180 |
| Number of negative individuals | 28 | 14 | 21 | 27 | 10 | 100 |
| Positivity (%) | 68 | 55 | 63 | 56 | 77 | 65 |
| Mean age | 5.7 | 4.5 | 4.2 | 6.5 | 5.0 | 5.5 |
| Area (m<sup>2</sup>) | 8500 | 3400 | 850 | 730 | 1200 | 14680 |
| Area per individuals (m<sup>2</sup>) | 97.7 | 109.7 | 14.9 | 12.2 | 27.3 | 52.6 |

We then investigated the cause of high STLV-1 prevalence. We hypothesized that a substantial proportion of infected macaques may play a critical role as viral super-spreaders for efficient inter-individual transmission, likely due to abnormally high PVLs and eventual incidence of poor humoral immune response against STLV-1. To examine this possibility, we evaluated ABTs and PVLs in the JM cohort and found that the ABTs among 180 seropositive macaques were normally distributed with a geometric mean of 4076 and an ABT of 8192 at the maximum number of individuals (Fig. 1A, Fig. S1). We observed no obvious differences in the titers between males and females (Fig. 1B) or among the five troops (Fig. 1C). We also examined the STLV-1 PVLs in the JMs PBMC samples and found that the PVLs among 168 macaques positive for the proviral DNA were normally distributed and ranged from 0.01%–20% with a geometric mean of 0.62% and PVLs of 0.64%–1.28% at the maximum number of individuals (Fig. 2A, Fig. S2). Again, we observed no statistical differences in the PVLs between males and females (Fig. 2B) or among the troops (Fig. 2C). The data regarding ABTs and PVLs from the 183 macaques positive for either value (herein tentatively regarded as ‘STLV-1-infected’) were plotted as shown in Figure 3. Among the JMs, 168 were positive for both values, whereas three were negative for ABTs but positive for PVLs, and 12 were positive for ABTs but negative for PVLs. Contrary to our expectations, we observed no monkeys with abnormally high PVLs and poor ABTs (Fig. 3). It is notable that the three ABT^-^ PVL^+^ monkeys belonged to two troops (two macaques in troop C and one in troop D), and their PVLs were comparable or less than the mean PVLs. It is, therefore, unlikely that only three monkeys caused the high prevalence in all the independent troops. In addition, we observed positive correlation between ABTs and PVLs (R = 0.50, *p* < 0.0001) (Fig. 3), suggesting that humoral immunity was properly induced in response to the increasing viral loads in these macaques.

**Figure 1.**
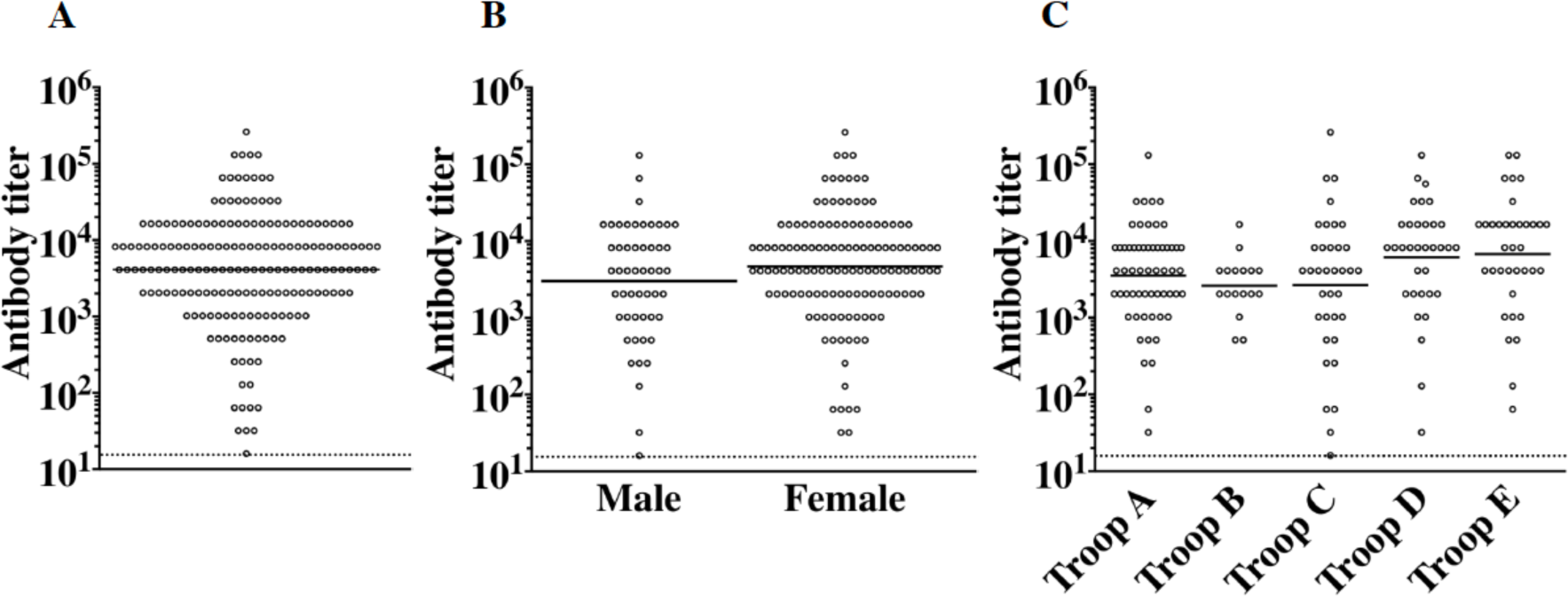
Distribution of anti-STLV-1 antibody titers (ABTs) in seropositive JMs. (A) Distribution of ABTs in all seropositive cohort JMs. (B) Results of the ABT distribution between male and female JMs and (C) among five troops are indicated. The dotted line shows the detection limit of the ABT, and the horizontal line indicates the geometric mean of the ABT distribution.

**Figure 2.**
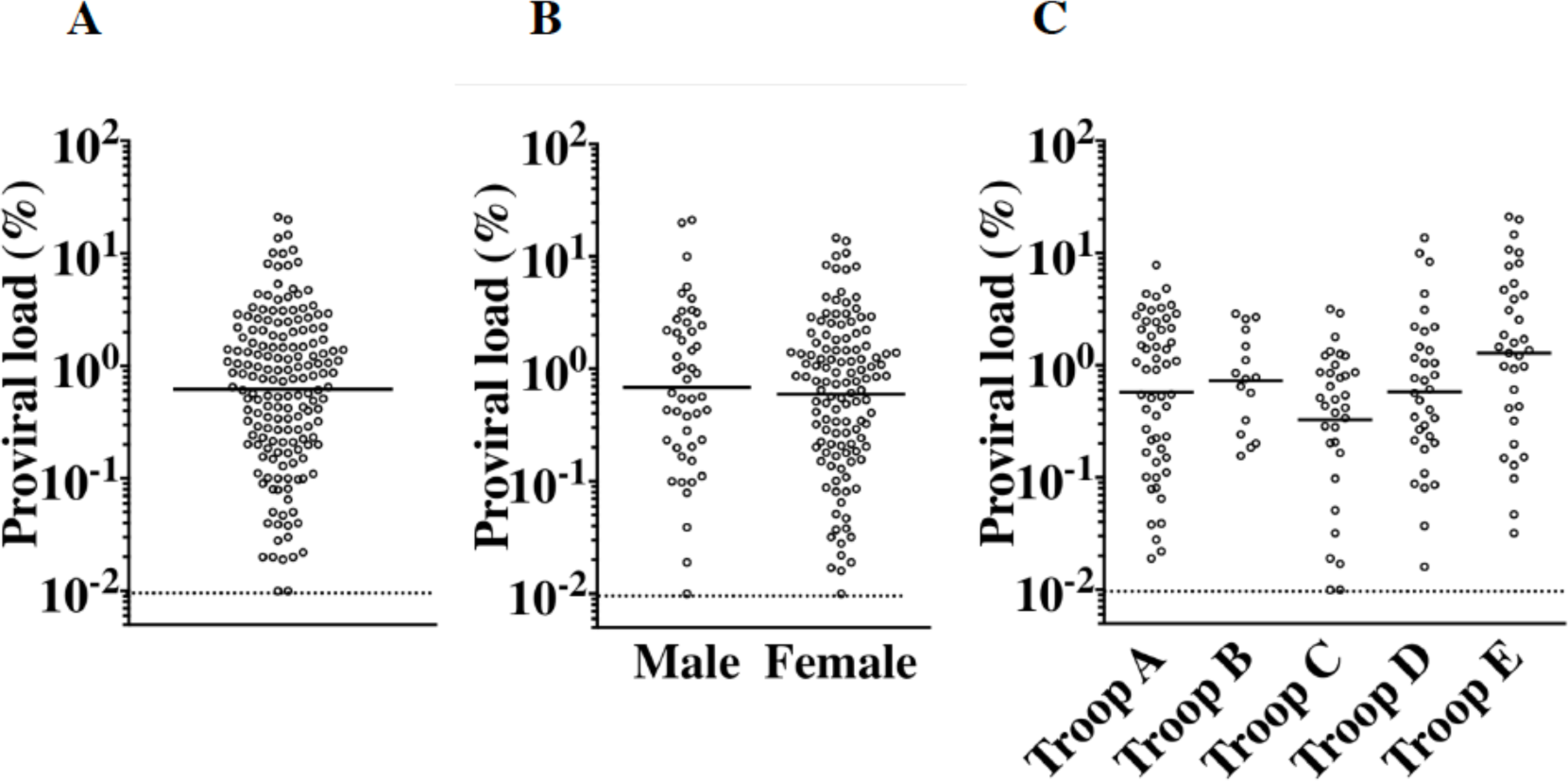
Distribution of proviral loads (PVLs). (A) Distribution of STLV-1 PVLs in proviral DNA-positive JMs. Results of the PVLs distribution between (B) male and female JMs and (C) among five troops (C) are shown. The dotted line indicates the detection limit of the PVL, and the horizontal line indicates the geometric mean of the PVL distribution.

**Figure 3.**
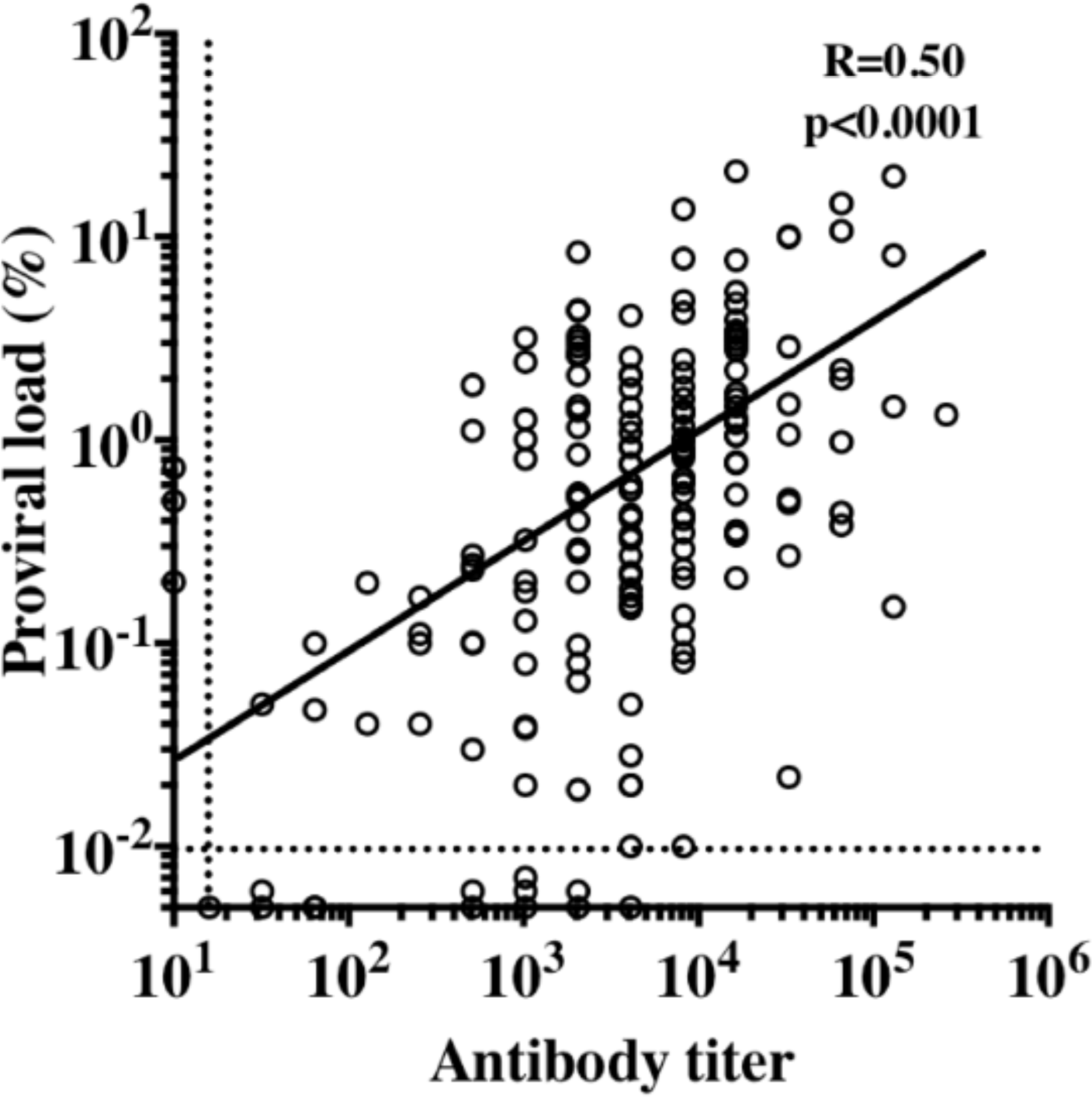
Correlation between antibody titers (ABTs) and proviral loads (PVLs) among individuals who were positive for either value. Among the macaques (N = 183), 168 were positive for both values, whereas three were seronegative but positive for PVLs, and 12 were seropositive but negative for PVLs. There was a significant correlation between the ABTs and the PVLs (R = 0.50; p < 0.0001).

In the absence of potential viral super-spreaders, we aimed to clarify the possible route(s) of transmission by which this high prevalence occurred. If maternal transmission were the main route of infection, the infection rate would drastically increase at around one year of age, followed by a gradual increase with age. On the other hand, if horizontal transmission were the main route, the infection rate would be low in younger ages, followed by a steep increase with age. To verify these possibilities, we examined the age-dependent change of seroprevalence in the cohort. The frequencies of seropositive individuals in each age group were 19%, 33%, 58%, 79%, 95%, 100%, and 96% at age groups of 0, 1, 2, 3–5, 6–9, 10–11, and ≥12 years, respectively (Fig. 4, solid line). We also analyzed the age-dependent change of proviral DNA prevalence (Fig. 5). The frequencies of proviral DNA-positive individuals in each age group were 13%, 33%, 55%, 75%, 91%, 100%, and 93% for age groups at 0, 1, 2, 3–5, 6–9, 10–11, and ≥12 years of age, respectively, which was consistent with those shown in Fig. 4. These results indicate that the infection rate drastically increased after one year of age and most of these animals over 6 years of age were infected with STLV-1, which supports the latter hypothesis that horizontal transmission would be the major route. Importantly, relatively large numbers of younger individuals (i.e., 0–1 years of age) whose STLV-1 prevalence was relatively low, apparently reduced the total prevalence to 65%. However, almost all of the adult individuals (i.e., sexually mature ones of more than 6 years of age) were infected with STLV-1 (Figs. 4 and 5, bar graphs). Each troop showed comparable results in both parameters (data not shown).

**Figure 4.**
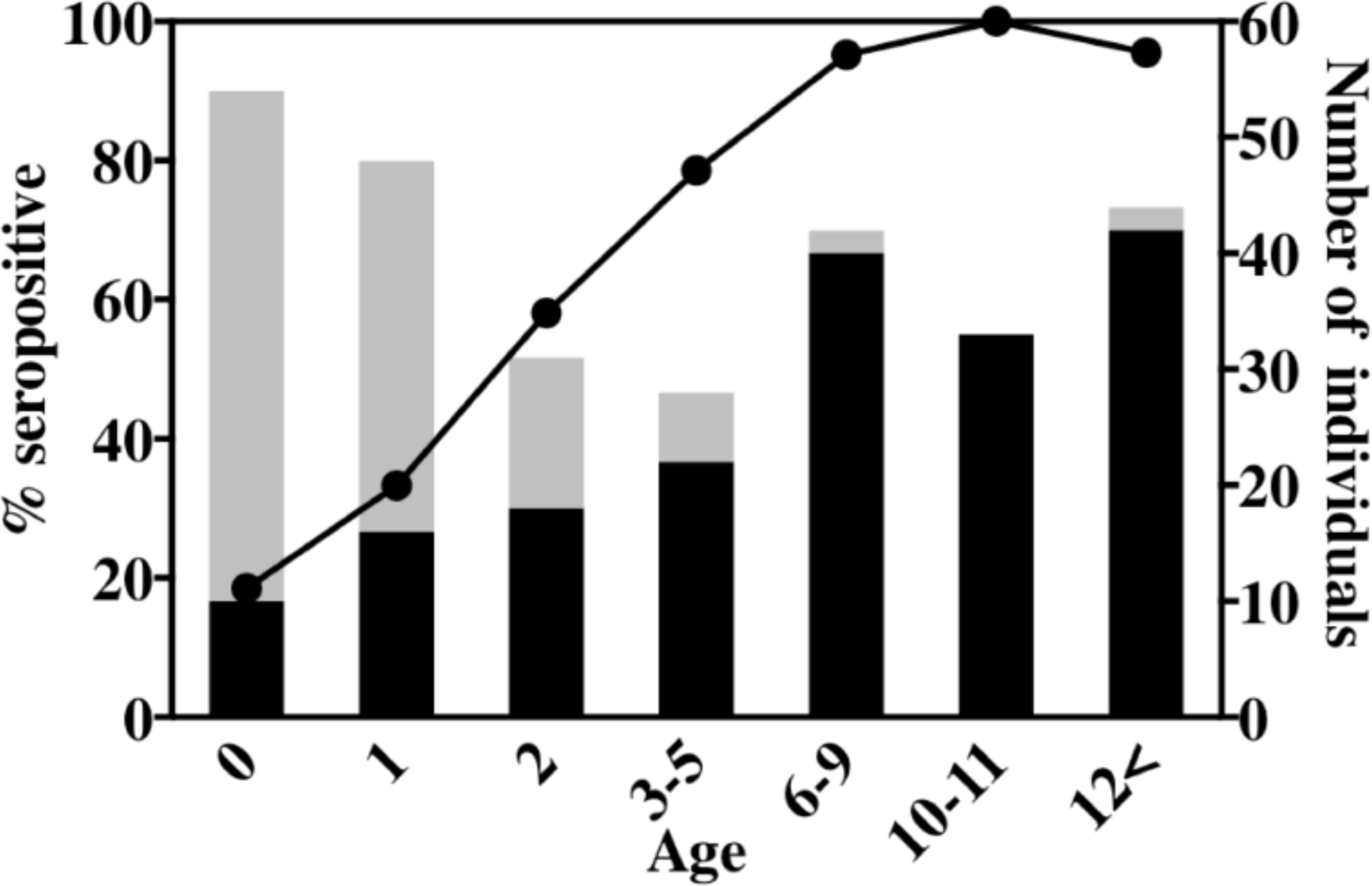
Age-dependent changes of STLV-1 seroprevalence in JMs. The left Y-axis shows the percentage of seropositive individuals (solid line). The right Y-axis indicates positive (closed bars) and negative (open bars) number of individuals.

**Figure 5.**
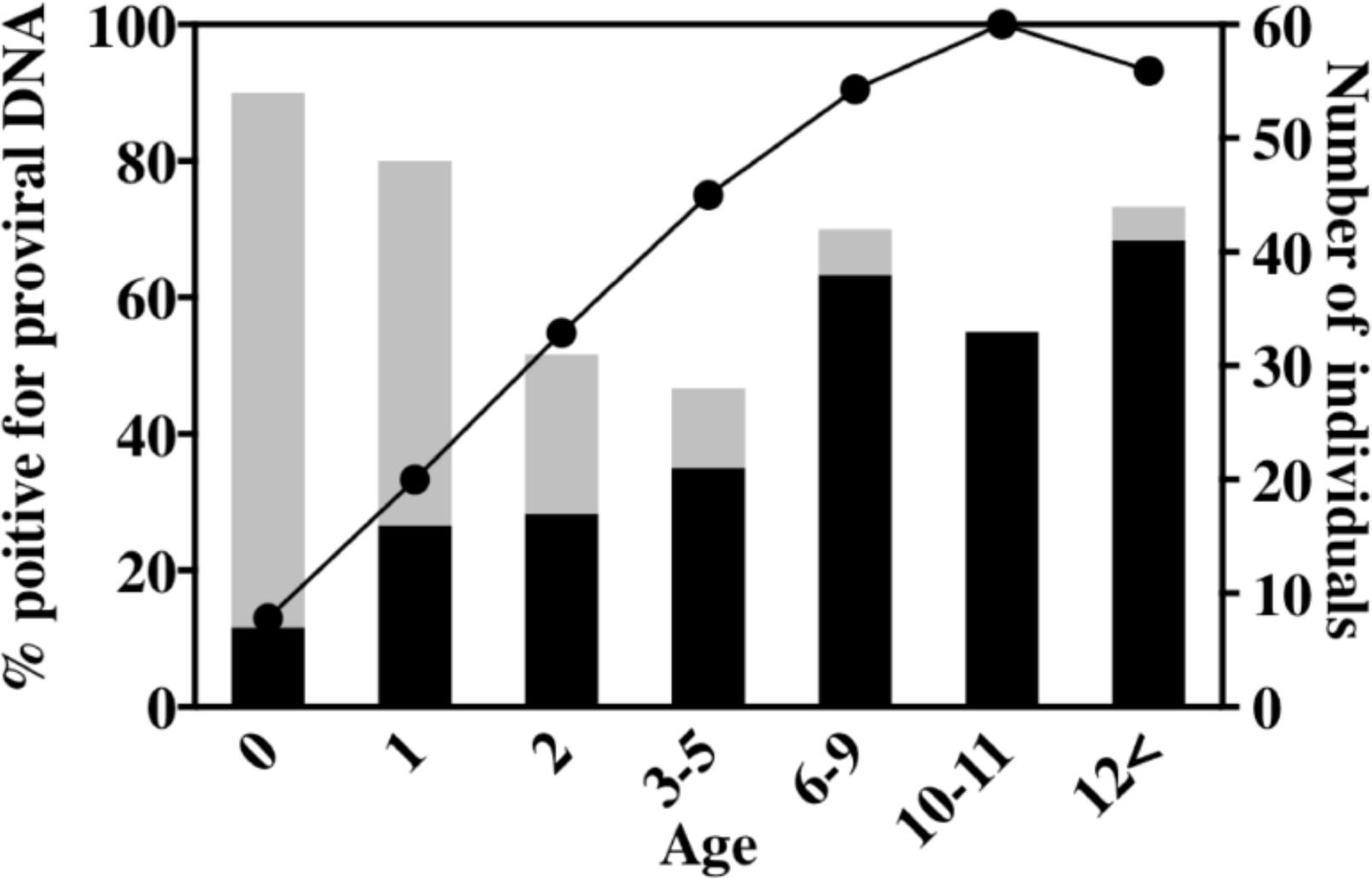
Age-dependent changes in the prevalence in JM positives for STLV-1 proviral DNA. The left Y-axis shows the percentage of proviral DNA-positive individuals (solid line). The right Y-axis indicates positive (closed bars) and negative (open bars) number of individuals.

Results described above suggest horizontal transmission as the major route of STLV-1 infection. There still remains a possibility that the seroconversion in the offspring of STLV-1-infected mothers, after the establishment of maternal transmission, could require up to three years due to long-term latency as shown in the case of HTLV-1 (65–67). If this is the case, then maternal transmission, rather than horizontal transmission, could be the major route. Therefore, we conducted a longitudinal study of the STLV-1 seroprevalence in this cohort (Table 2). We selected 139 monkeys whose serum samples in both 2011 and 2015 were available (PBMC samples in 2011 were not available). In 2011, 111 of 139 monkeys were seropositive, whereas 28 were seronegative. It was found that among the 28 seronegative monkeys in 2011, 24 were seroconverted for the antibody within four years from 2011 to 2015. Remarkably, among ten seronegative monkeys of four years old and above in 2011, eight were seroconverted within four years (80%), which was comparable with the monkeys of three years old and below in 2011 (16/18, 89%). The fact that frequent seroconversion occurred even in the seronegative monkeys of four years old and above suggests lower probability of long-term latency post-maternal transmission and supports the notion that horizontal STLV-1 transmission frequently occurs among JMs, which may eventually result in almost all adult monkeys infected with STLV-1.

**Table 2:** Longitudinal study of the STLV-1 prevalence in Japanese macaques.

| Age in 2011<br>(Age in 2015) | Number of<br>seronegative<br>in 2011 | Number of<br>seropositive in<br>2015 | Frequency of<br>seroconversion |
| --- | --- | --- | --- |
| 0 (4) | 7 | 6 |  |
| 1 (5) | 1 | 1 |  |
| 2 (6) | 6 | 6 |  |
| 3 (7) | 4 | 3 |  |
| 0-3 (4-7) | 18 | 16 | 16/18 (89%) |
| 4 (8) | 3 | 3 |  |
| 5 (9) | 1 | 1 |  |
| 6 (10) | 3 | 3 |  |
| ≥7(≥11) | 3 | 1 |  |
| ≥4(≥8) | 10 | 8 | 8/10 (80%) |
| Total | 28 | 24 | 24/28 (86%) |

## Discussion

In this study, we aimed to investigate the cause of the high prevalence of STLV-1 infection in the studied JMs cohort. We initially examined the prevalence of STLV-1 infections in the JMs derived from five independent troops originating from inhabitants of different areas and found that 65% (180/280) of the macaques were seropositive, which was generally consistent with previous reports (47, 48, 50, 52) (Table 1). Contrary to our expectations, we found that the ABTs and PVLs among the infected macaques were normally distributed with mean values of 4076 and 0.62%, respectively (Figs. 1, 2, S1, and S2). This was comparable to those of HTLV-1-infected humans. In addition, we did not observe macaques with abnormally high PVLs and poor ABTs (Fig. 3). Thus, the possibility of viral super-spreaders is unlikely. To further determine the possible route(s) of transmission, the influence of age on frequency of STLV-1 infection in the cohort was examined. We found that the frequency drastically increased over one year of age and most of these animals over 6 years of age were infected with STLV-1 (Figs. 4 and 5). Moreover, the longitudinal follow-up study of this cohort demonstrated that frequent seroconversion occurred in not only infants but also in juvenile and adult seronegative monkeys (Table 2). Taken together, our findings strongly suggest that frequent horizontal transmission is the major route of STLV-1 infection in JMs, which eventually result in almost all adult monkeys infected with STLV-1. These findings were unexpected considering human cases of HTLV-1 infection, of which the prevalence rate is only 1% (or below) in Japan (an endemic country) (16). What causes the high frequency of horizontal STLV-1 transmission in JMs? It was shown that JMs genetically originate from rhesus macaques (RMs) as the ancestor macaques came over from the Asian Continent to Japan around 0.5 million years ago (68). It was reported that much less frequency of RMs are infected with STLV-1 than the case of JMs (69). Similarly, the prevalence rate of STLV-1 in RMs bred and reared in our free-ranging facility as well as JMs is less than 1% (52). It is therefore reasonable to speculate that STLV-1 was broadly disseminated after ancestor macaques started inhabiting Japan. As for the migrated JMs, foods such as leaves, fruits, and nuts in their habitats were insufficient in the cold winter season so they probably needed to form troops to keep their territories for foods and to stay warm by assembling together (70). They eventually established a promiscuous mating system without having fixed partners/mates to circumvent the genetic disadvantages caused by inbreeding within the troop (71). It is possible that promiscuity increased the opportunity to transmit STLV-1, which led to the high STLV-1 prevalence. In fact, it was reported that a relatively high prevalence of HTLV-1 was occasionally observed in isolated Japanese populations (72), which is generally consistent with the phenomenon observed in JMs.

Results obtained in this study indicate that less than 20% of infants (i.e., 0-year-old) were positive for either antiviral antibodies or proviral DNA (Figs. 4 and 5). This is generally comparable with the estimated frequency of maternal transmission of HTLV-1 in humans (73). However, it remains to be elucidated whether long-term latent STLV-1 infection in infants and eventual seroconversion from latency of a couple of years after birth could occur frequently. It was shown that frequency of maternal transmission was associated with the PVLs of the pregnant mothers (74–77). If this is the case in JMs, this suggests that mean PVLs, as well as their distribution among the macaques (Figs. 1 and 2), are similar to human cases (78, 79), and this may support the possibility that frequency of maternal STLV-1 transmission might be comparable to humans. It is intriguing to determine the frequency of mother-to-child STLV-1 transmission as well as the period of time required for the seroconversion in the mother-to-child transmission as done herein.

## Materials and methods

### Animals

JMs bred and reared in the free-range facility of the Primate Research Institute, Kyoto University (KUPRI) were used in this study. All the troops were isolated and had no physical connection with each other. All animal experiments were approved by the Animal Welfare and Animal Care Committee of KUPRI (approval numbers: 2014–092, 2015-040, and 2016-135) and were conducted in accordance with the Guidelines for Care and Use of Nonhuman Primates (Version 3) by the Animal Welfare and Animal Care Committee of KUPRI.

### Preparation of plasma and peripheral blood mononuclear cells (PBMCs)

Blood samples were collected from JMs at routine health checkups under ketamine anesthesia with medetomidine, followed by administration of its antagonist, atipamezole, at the end of the procedure. PBMCs were separated from blood samples with Ficoll-paque PLUS (GE Healthcare, Buckinghamshire, UK) by density gradient centrifugation. Plasma and PBMCs were frozen at −80°C until use. Cellular DNA was purified via a QIAamp DNA Blood Mini Kit (Qiagen, Hilden, Germany), according to the manufacturer’s instructions.

### Titration of the STLV-1-specific antibody

Plasma samples were evaluated for ABTs with a particle-agglutination assay using Serodia-HTLV-1 (Fujirebio Inc. Tokyo, Japan) as previously described (52). The plasma cut-off titer was a 1:16 dilution.

### Quantification of STLV-1 PVLs

Cellular DNA collected from PBMCs was measured for STLV-1 PVLs via a real-time PCR quantification of copy numbers of the STLV-1 *tax* gene and *RAG1* gene of JMs as previously described (52). PCR was performed using Thunderbird Probe qPCR mix (TOYOBO, Osaka, Japan). The following primers and probes were used: RAG1-2F (CCCACCTTGGGACTCAGTTCT), RAG1-2R (CACCCGGAACAGCTTAAATTTC), a RAG1 probe (5’-FAM CCCCAGATGAAATTCAGCACCCATATA TAMRA-3’), STLV-1 tax-F2 (CTACCCTATTCCAGCCCACTAG), STLV-1 tax-R3 (CGTGCCATCGGTAAATGTCC), and a STLV-1 tax probe (5’-FAM CACCCGCCACGCTGACAGCCTGGCAA TAMRA-3’). Copy number of STLV-1 proviral DNA per cell was standardized with that of the *RAG1* gene. The detection limit of PVLs was 0.01%.

### Statistical analyses

We tested the normal distribution of the data and applied parametric or non-parametric methods according to the experiment. Pearson’s correlation coefficient was employed for correlation of two parameters, and two-tailed Student’s *t*-tests were employed for comparison of two groups. For multiple comparisons with more than two groups, a one-way ANOVA with Tukey’s multiple comparison test was used.

## Acknowledgements

This research was supported by AMED under Grant Number JP19fk0108059. The blood samples of JMs used in this study were partly provided by National Bio-Resource Project “Japanese Monkey” of MEXT, Japan. We thank Kaoru Tsuji and colleagues of Center for Human Evolution Modeling Research in KUPRI for their technical assistance.

## Supplementary Information

**Figure S1.**
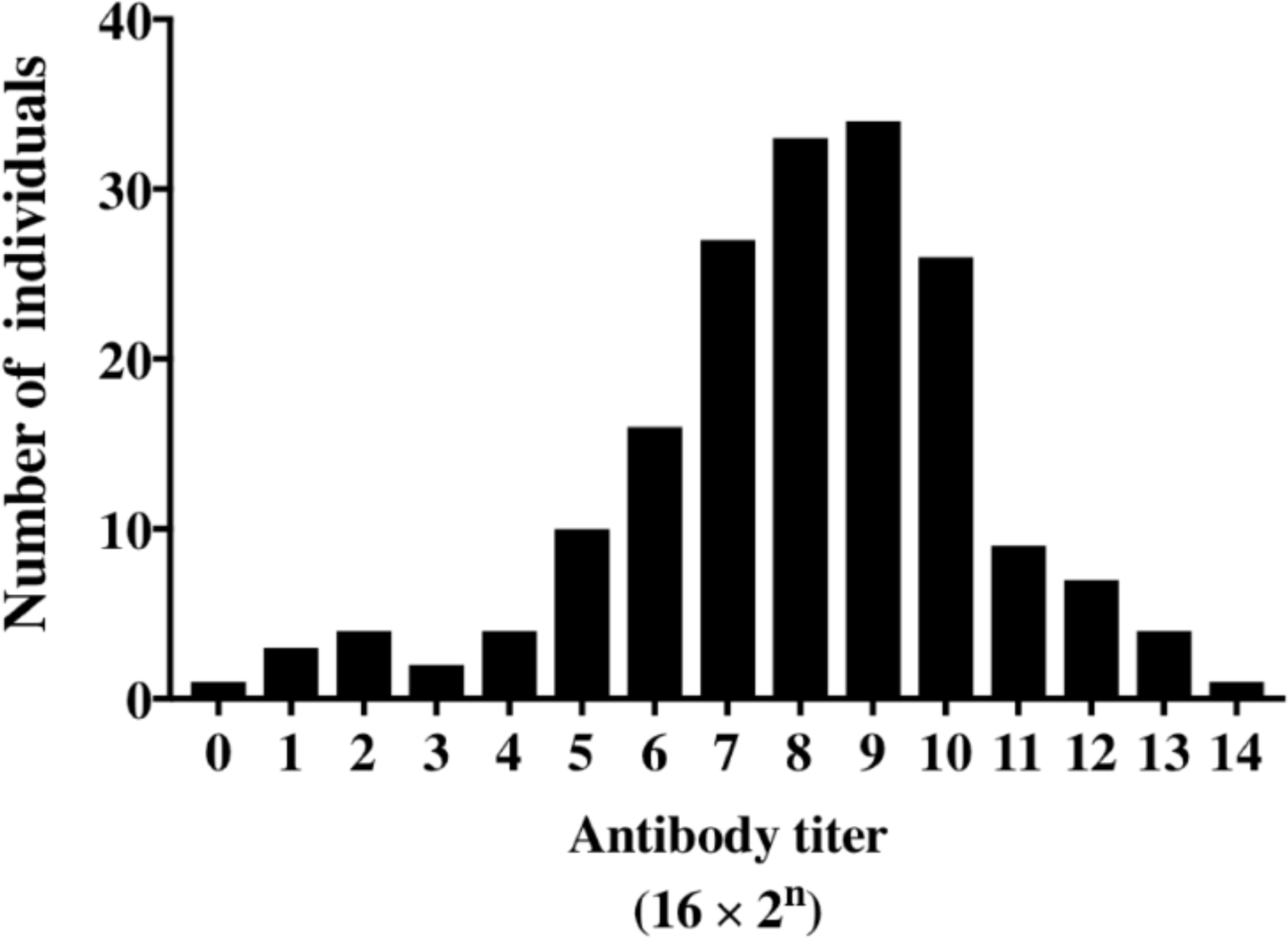
Distribution of anti-STLV-1 antibody titers (ABTs) in seropositive JMs. The X-axis represents antibody titers ranging from 16–262144, with an ABT of 8192 at the maximum number of individuals. The Y-axis represents the number of individuals in each antibody titer.

**Figure S2.**
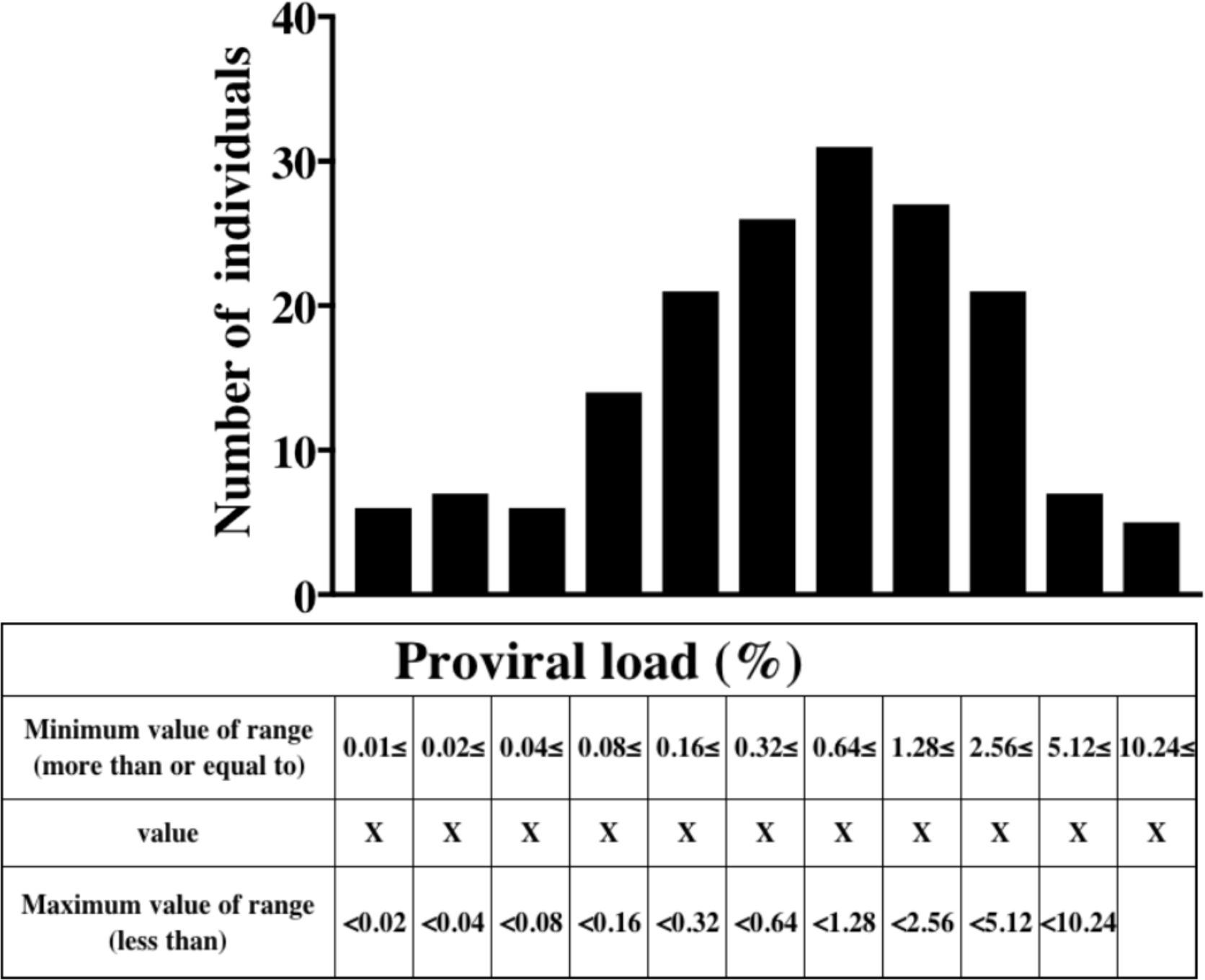
Distribution of STLV-1 proviral loads (PVLs) Distribution of PVLs in JMs positives for STLV-1 proviral DNA. The X-axis indicates PVLs ranging from 0.01%–20%, with PVLs of 0.64%–1.28% at the maximum number of individuals. The Y-axis shows the number of individuals in each PVL group.

